# Long-acting hydrogel-based depot formulations of tirzepatide and semaglutide for the management of type 2 diabetes and weight

**DOI:** 10.1101/2025.07.02.662867

**Authors:** Andrea I. d’Aquino, Changxin Dong, Leslee T. Nguyen, Jerry Yan, Carolyn K. Jons, Olivia M. Saouaf, Ye Eun Song, Noah Eckman, Sara Kapasi, Christian M. Williams, Vannessa Doulames, Samya Sen, Manoj K. Manna, Alakesh Alakesh, Katie Lu, Ian Hall, Eric A. Appel

## Abstract

Several incretin hormone therapies have been clinically approved and have revolutionized the treatment of diabetes and obesity. Promising therapeutics include semaglutide (Ozempic^®^ and Wegovy^®^), an agonist for glucagon-like peptide-1 (GLP-1) receptor, and tirzepatide (Mounjaro^®^), a dual agonist for GLP-1 and glucose-dependent insulinotropic polypeptide (GIP) receptors. These molecules help to regulate blood glucose levels, enhance insulin secretion and sensitivity, and reduce appetite. Currently, these treatments require weekly injections, which can be challenging for patients to adhere to. We recently reported the development of an injectable hydrogel depot technology enabling months-long release of semaglutide (Sema). Here, we further develop this technology for improved prolonged release of both Sema and tirzepatide (TZP). In a rat model of diabetes, we show a single administration of hydrogel-based formulations of either Sema or TZP maintained relevant drug levels for over 6 weeks. In these studies, single administrations of long-acting hydrogel-based therapies of Sema or TZP were similarly effective at regulating blood glucose and weight compared to daily injections of either Sema or TZP in standard aqueous vehicles. This hydrogel depot technology is easy to manufacture, injectable, and exhibits excellent biocompatibility, enabling months-long-acting treatments with the potential to improve management of diabetes and weight.

## 1. Introduction

Diabetes affects an estimated 530 million adults globally, ^[1, 2]^ with type 2 diabetes (T2D) comprising over 98% of all diagnosed cases.^[3–5]^ T2D is a progressive metabolic disorder characterized by insulin resistance, impaired insulin secretion, obesity, diminished incretin response, dyslipidemia,^[3, 6–9]^ and ultimately in β-cell dysfunction.^[10]^ Recent therapeutics strategies have prioritized incretin-based therapies, notably glucagon-like peptide-1 receptor agonists (GLP-1 RAs) such as semaglutide (Ozempic^®^, Wegovy^®^) and liraglutide (Victoza^®^), which improve glycemic control, reduce glycated hemoglobin (HbA1c), and promote weight loss. ^[11]^ Incretins, primarily GLP-1 and gastric inhibitory polypeptide (GIP), stimulate insulin secretion in a glucose-dependent manner and modulate satiety, energy intake, and lipid metabolism. Incretin hormones, gastric inhibitory polypeptide (GIP) and glucagon-like peptide-1 (GLP-1), both play important roles in triggering the pancreas to secrete insulin to achieve normal glucose homeostasis, however, for people with diabetes, the pancreas becomes less responsive to incretins (predominately to GIP).^[11]^ Over the past decade, pharmacological-level replacement of GLP-1 receptor agonism has become one of the most effective tools for improving glucose homeostasis, weight loss and cardiovascular risk in patients with diabetes.^[11, 12]^ While GLP-1 has long been used pharmacologically, combining GIP and GLP-1 receptor agonism, as in tirzepatide (Mounjaro^®^), further improves insulin sensitivity and weight loss outcomes compared to GLP-1 RA monotherapy.^[13, 14]^ In patients with T2D, GIP counteracts insulin-induced hypoglycemia, most likely through a glucagonotropic effect.^[13]^ In contrast, during hyperglycemia, GIP increases glucose disposal through a predominant effect on insulin release.^[13]^ Additionally, GIP amplifies adipose-tissue sensitivity to insulin, facilitating the role of adipose tissue in lipid buffering after meals and thus preventing the deposition of fat outside of traditional fat deposits^[6]^ and it appears to play a role in enhancing GLP-1-mediated central satiety.^[14]^ For both GIP and GLP-1, beneficial effects on cardiovascular health have been observed, and recent studies have shown that GIP/GLP-1 receptor co-agonists have superior efficacy compared to selective GLP-1 receptor agonists with respect to glycemic control as well as body weight.^[14]^ (**Figure 1A**)

**Figure 1.**
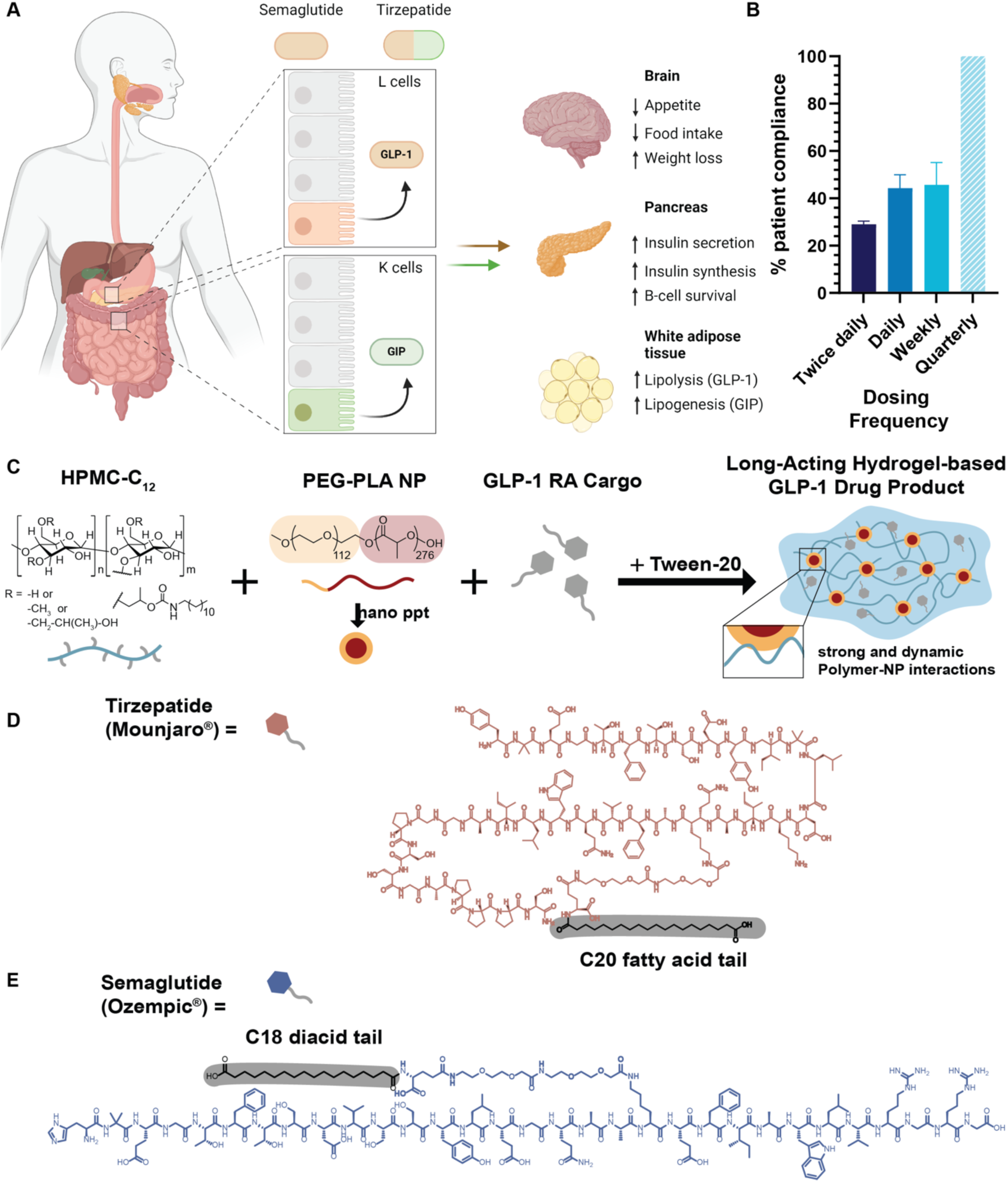
PNP hydrogels enable prolonged delivery of GLP-1 receptor agonists. **(A)** Schematic of the biological actions of glucose-dependent insulinotropic polypeptide (GIP) and glucagon-like peptide-1 (GLP-1) at the organ and tissue level. **(B)** Reported patient adherence rates show limited improvement with once-weekly dosing compared to once-daily regimens; a once-quarterly dosing schedule (striped bar) represents a hypothetical ideal for maximizing compliance. **(C)** Schematic of PNP hydrogel formulation, composed of hydrophobically modified HPMC-C12, PEG-PLA nanoparticles, and polysorbate 20 (Tween-20), enabling encapsulation of GLP-1 receptor agonists such as tirzepatide (TZP) or semaglutide (Sema). **(D)** Chemical structure of tirzepatide. **(E)** Chemical structure of semaglutide.

Approved in 2023, tirzepatide (TZP) is a dual GIP/GLP-1 receptor agonist that significantly enhances glycemic control and is under clinical investigation for obesity, cardiovascular disease, and non-alcoholic steatohepatitis.^[6]^ It is also being studied for its potential in chronic weight management, reducing major adverse cardiovascular events, amongst other conditions such as heart failure, obesity, and non-cirrhotic non-alcoholic steatohepatitis.^[6]^ Currently, these drugs necessitate frequent administration, with dosages ranging from daily to weekly (twice daily and weekly exenatide; daily liraglutide; weekly dulaglutide; and weekly semaglutide).^[12]^ High costs and the demanding frequency of drug administration leads to adherence rates of less than 50%, with these rates declining further over time.^[15–17]^ (**Figure 1B**) Nonadherence is responsible for 10% to 25% of hospital admissions and increases the risk of hospitalization by 2.5 times in diabetic patients.^[18, 19]^ This issue has led to an estimated $100 to $300 billion in healthcare costs each year, which could be reduced by improving medication adherence.^[17, 18]^ Strategies including simplifying treatment regimens, reducing side effects, and enhancing drug delivery methods can lead to better patient outcomes.^[20]^

Hydrogels offer a promising platform to address these challenges by enabling sustained, controlled delivery of peptide drugs with favorable biocompatibility and depot persistence.^[21, 22]^ In particular, we have previously developed polymer–nanoparticle (PNP) hydrogels formed via self-assembly of biodegradable PEG-PLA nanoparticles and hydrophobically modified hydroxypropyl methylcellulose (HPMC-C12).^[23–30]^ These supramolecular hydrogels exhibit injectability, self-healing, and tunable rheological properties, supporting their use as long-acting delivery vehicles for GLP-1 RAs. PNP hydrogels are a versatile platform that has demonstrated extended release profiles for diverse therapeutic modalities, such as vaccines,^[31, 32]^ semaglutide and liraglutide,^[33]^ antibodies,^[34]^ cancer therapies,^[35]^ and even serving as environmental protection agents such as wildfire retardants.^[36, 37]^ Such vehicle serves as a promising delivery strategy for sustained GIP/GLP-1 receptor co-agonists.

In this study, we expand the application of PNP hydrogels Sema depots^[38]^ to formulate long-acting TZP depots, engineered for months-long, steady-state release following a single injection (**Figure 1C**–**E**), and compare their therapeutics effects and pharmacokinetics release profile. The hydrogel network, which preserves the native aqueous environment of peptide drugs, circumvents common formulation challenges of other depot technologies and aligns with the regulatory requirements for aqueous peptide-based products. By leveraging the favorable material properties of PNP hydrogels, this work aims to provide a next-generation injectable therapeutic platform to transform T2D management and improve adherence across chronic treatment landscapes.

## 2. Results

### 2.1. Design of injectable hydrogels for sustained release of TZP

TZP has demonstrated clinically significant benefits in reducing HbA1c, improving glycemic control, and promoting weight loss in adults with T2D.^[6]^ While both Sema and TZP are effective GLP-1 receptor agonists, once-weekly TZP has shown superior clinical efficacy compared to Sema, making it a promising therapeutic for T2D management.^[39, 40]^ Structurally, TZP contains a pendant C20 fatty diacid moiety that promotes albumin binding and extends systemic circulation. We hypothesized that this lipophilic feature could facilitate non-covalent incorporation of TZP into the hydrophobic domains of the PNP hydrogel network during formulation, without requiring chemical modification (**Figure 1D**). This interaction is enabled by the hydrophobic nature of both components in the hydrogel, including the poly(lactic acid) (PLA) core of the PEG-PLA nanoparticles and the dodecyl moieties on the HPMC-C12 polymers. These features provide multiple binding sites for hydrophobic interactions, facilitating physical entrapment of the peptide within the hydrogel matrix.^[33]^ Based on the clinical dosing regimen (2.5 to 5 mg TZP weekly), we estimated that a depot designed to release TZP continuously over four months in a rat model should contain approximately 4.5 mg of drug within a subcutaneously injectable volume (typically 0.5 to 2 mL).

To formulate the PNP hydrogels, a suspension of biodegradable PEG-PLA nanoparticles containing the therapeutic peptide was mixed with a solution of dodecyl-modified hydroxypropyl methylcellulose (HPMC-C12) (**Figure 1C**).^[28]^ To prevent peptide aggregation during formulation, a minimal amount of the surfactant polysorbate 20 (Tween-20) was added. Tween-20 has been widely reported to suppress aggregation of peptides and proteins by stabilizing them in solution, and its inclusion here supports the maintenance of drug integrity throughout hydrogel fabrication and storage. ^[33, 41, 42]^

In practice, the PEG-PLA nanoparticle solution (containing the peptide and Tween-20) is loaded into one sterile syringe, while the HPMC-C12 solution is loaded into a second. The two components are then mixed via a sterile Luer-lock elbow connector, initiating rapid, multivalent hydrophobic interactions between the nanoparticles and polymer chains. This results in physical cross-linking and formation of a robust, solid-like hydrogel (**Figure 1C**). Despite their solid-like properties, the hydrogels remain shear-thinning and can be delivered through narrow-bore needles using clinically practical injection forces. As demonstrated in syringe pump tests, injection forces for these formulations remain within 0–5 N at flow rates of 0.5–2 mL min^−1^ (**Figure 2A**). Importantly, the viscoelastic and drug release characteristics of the hydrogel can be precisely tuned by altering the relative concentrations of HPMC-C12 and PEG-PLA nanoparticles. This tunability enables the design of customized depot systems tailored for specific release kinetics and clinical needs. Previous studies have shown that PNP hydrogels are highly biocompatible, exhibit minimal swelling under physiological conditions, and are broadly suitable for biomedical use. ^[21, 25, 33]^ Taken together, these properties position the PNP hydrogel system as a modular, injectable platform capable of supporting sustained, long-acting delivery of complex biologics such as TZP.

**Figure 2.**
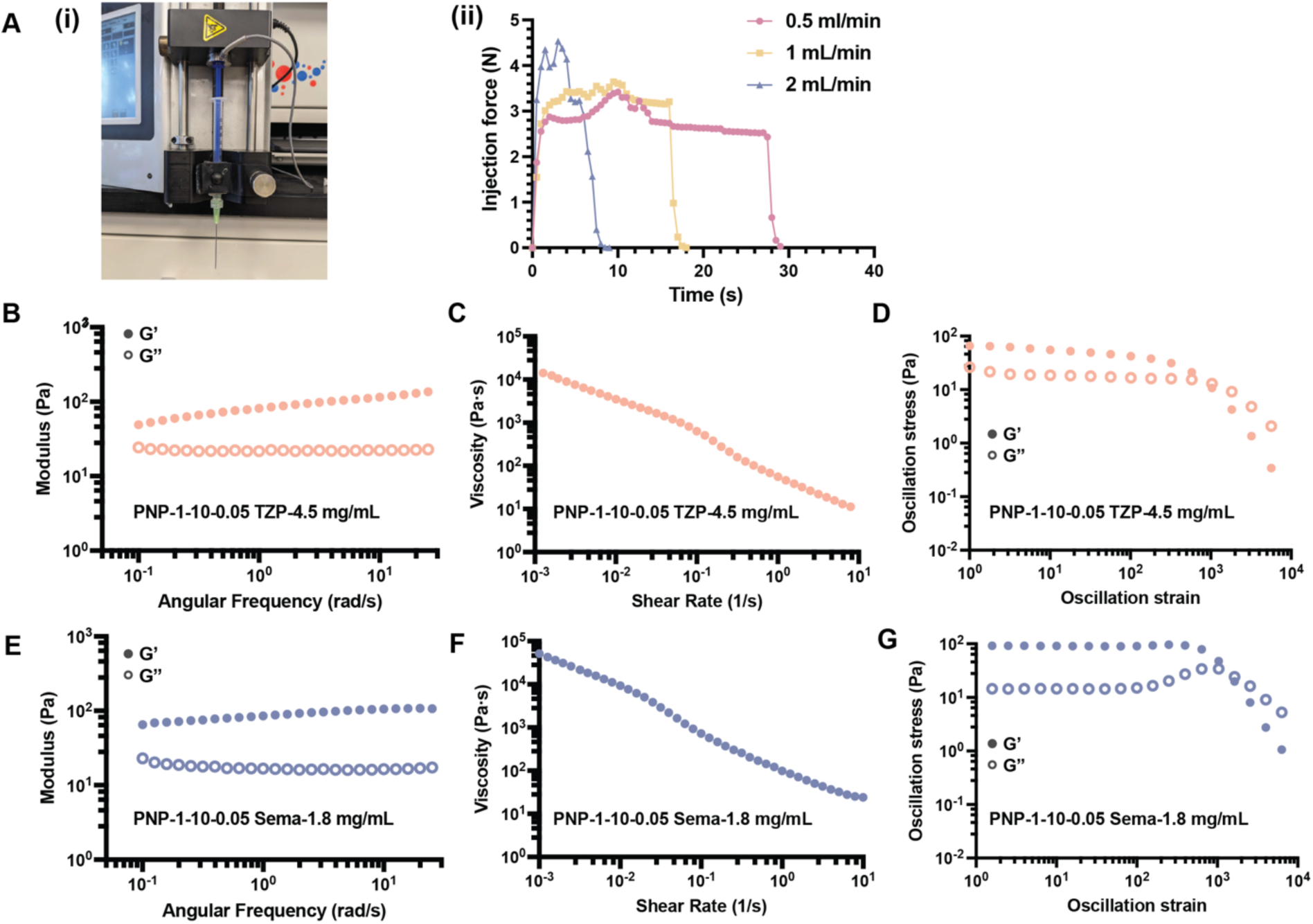
Mechanical properties of PNP hydrogels loaded with GLP-1 receptor agonists. PNP hydrogels are formed by mixing hydrophobically modified HPMC (HPMC-C12) with PEG-PLA nanoparticles and GLP-1 cargo using a Luer lock mixer, yielding a homogeneous, bubble-free hydrogel. Owing to dynamic physical crosslinking, these hydrogels are injectable through high-gauge needles and self-heal after injection. **(A)** Injection force was evaluated using **(i)** a 40 mm 21G needle setup and **(ii)** recorded over a 30 s injection period. **(B)** Rheological characterization of TZP-loaded PNP-1-10-0.05 hydrogels: **(i)** frequency-dependent oscillatory shear sweep, **(ii)** flow curve illustrating shear-thinning behavior, and **(iii)** strain sweep showing the linear viscoelastic region. **(C)** Rheological characterization of Sema-loaded PNP-1-10-0.05 hydrogels, showing similar trends across **(i–iii)**.

### 2.2. Rheological Characterization of GLP-1 PNP Hydrogels

To evaluate the impact of different GLP-1 receptor agonists on the mechanical behavior of PNP hydrogels, we prepared two formulations: **PNP-1-10-0.05 TZP-4.5 mg/mL** and **PNP-1-10-0.05 Sema-1.8 mg/mL**. In this nomenclature, the numbers denote the weight percentages (wt%) of polymer (HPMC-C12), nanoparticles (PEG-PLA), and Tween 20 (surfactant), respectively, while the remainder is buffer (phosphate buffered saline or PBS). The formulations were designed to deliver therapeutically relevant doses for translational evaluation in a rat model of insulin-impaired diabetes.

To determine whether the identity of the peptide cargo influences hydrogel mechanical or transport properties, such as cargo release behavior, we conducted rheological characterization of both formulations. These studies ensured that the hydrogels met criteria for *injectability*, subcutaneous *depot* formation, and long-term *stability* (**Figure 2B–D**, **E–G**). Frequency-dependent oscillatory shear testing demonstrated that both formulations exhibited predominantly solid-like viscoelastic behavior, with storage modulus (*G′*) exceeding loss modulus (*G′′*) over all observed frequencies, and tan(*δ*) values between 0.1 and 0.5 across the entire frequency range, indicative of robust physical cross-linking and structural integrity essential for depot formation (**Figure 2B**, **E**). In stress-controlled flow sweep tests, both samples displayed strong shear-thinning behavior and no evidence of fracture, enabling injection through small-diameter, clinically relevant needles (**Figure 2C**, **F**).^[43]^ The injectability of the formulations was verified using injection force measurements (**Figure A-i**, where the maximum force required to extrude the gels was below 5 N at relevant extrusion flow rates for both samples (**Figure A-ii**). Stress-controlled oscillatory shear rheology showed that both PNP hydrogels resisted flow below a certain stress or strain value and began to flow with a sharp decrease in shear stress once this threshold was exceeded (**Figure 2D, G**). This observation is in corroboration with flow sweep data and shows that the PNP hydrogels can smoothly transition from stiff, solid-like materials (linear regime at low strains) to shear-thinning fluids (nonlinear regime at large strains). The observed yield stress from flow sweeps (**Figure 2C, F**; *τ*y > 100 Pa) is critical for maintaining a stable subcutaneous depot following administration.^[21]^ Importantly, rheological comparisons between PNP-1-10-0.05 TZP-4.5 mg/mL and PNP-1-10-0.05 Sema-1.8 mg/mL (**Figure 2B–D** vs. **2E–G**) revealed no substantial differences in mechanical performance. This observation suggests that the identity of the encapsulated GLP-1 receptor agonist, TZP or Sema, does not significantly alter the bulk mechanical properties of the PNP hydrogel platform.

### 2.3. In vitro release kinetics of TZP from hydrogel depot

In vitro release profiles of TZP and Sema from PNP hydrogels were examined to assess how differences in peptide cargo might influence diffusion and retention. To enable a direct comparison, both PNP-1-10-0.05 TZP-1.8 mg/mL and PNP-1-10-0.05 Sema-1.8 mg/mL hydrogels were loaded into glass capillaries and overlaid with PBS under infinite sink conditions at 37 °C. The low injection volume allowed precise loading of 1.8 mg/mL of each peptide in the hydrogel, and aliquots were collected over time for analysis via high-performance liquid chromatography–mass spectrometry (HPLC–MS) (**Figure 3A**, **Method S1**). Over the course of three weeks, both formulations showed minimal release and reached a plateau after approximately one week, indicating rapid formation of a diffusion-limiting depot. The PNP-1-10-0.05 TZP-1.8 mg/mL formulation released 0.41% of its total payload, while the PNP-1-10-0.05 Sema-1.8 mg/mL formulation released 1.09% over the same time frame (**Figure 3B–C**). These results suggest that both peptides remain largely entrapped within the hydrogel matrix under passive release conditions. The low cumulative release observed in both cases reflects the tight physical network of the PNP hydrogel and the compatibility of both peptide cargos with the formulation. However, Sema exhibited slightly higher release than TZP, which may reflect differences in hydrophobicity, molecular conformation, or self-association behavior between the two peptides. Importantly, this distinction did not correspond to any measurable differences in rheological properties of the hydrogels (**Figure 2**), suggesting that the PNP platform provides a stable and tunable delivery scaffold that is largely agnostic to peptide cargo. Overall, these results demonstrate that both TZP and Sema can be effectively retained in PNP hydrogel depots with minimal passive diffusion, supporting the platform’s utility for long-acting subcutaneous delivery. Minor differences in release kinetics may be further optimized through formulation tuning or peptide-specific excipients.

**Figure 3.**
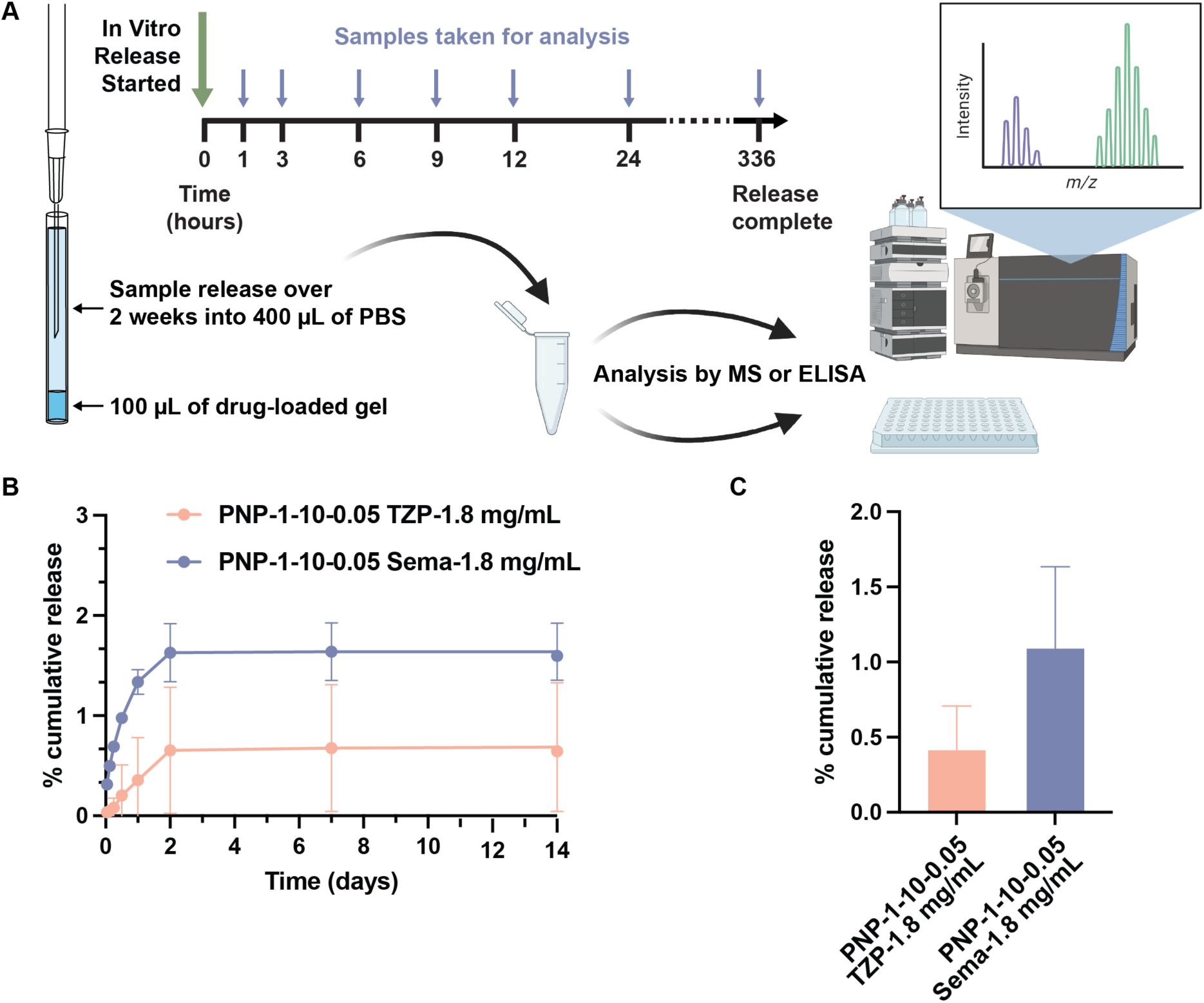
In vitro release of GLP-1 receptor agonists from PNP hydrogels. **(A)** Schematic of the in vitro release assay evaluating TZP and Sema release from PNP hydrogels over three weeks. Hydrogels were loaded into capillary tubes to minimize erosion by reducing the surface area-to-volume ratio. TZP concentrations were quantified by HPLC-MS, and Sema by ELISA. **(B)** Cumulative in vitro release profiles of TZP and Sema from PNP-1-10-0.05 hydrogel formulations over 21 days (n = 3). **(C)** Total cumulative release of each cargo from the hydrogel formulations at study endpoint, presented as bar graphs.

### 2.4. Evaluation of in-vivo pharmacokinetics and pharmacodynamics of hydrogel-based TZP formulations

Prolonged and controlled administration of TZP could provide a long-term treatment solution for type 2 diabetes, maintaining stable incretin hormone levels for several months instead of days or weeks. To assess the effectiveness of our hydrogel formulations in prolonging the delivery of TZP, we performed pharmacokinetic and pharmacodynamic studies using a rat model of insulin impairment characteristic of type 2 diabetes (**Figure 4A**). Diabetes was induced using nicotinamide (NA) and streptozotocin (STZ) in rats, an effective mimic of a T2D-like phenotype (**Method S2**).^[44, 45]^

**Figure 4.**
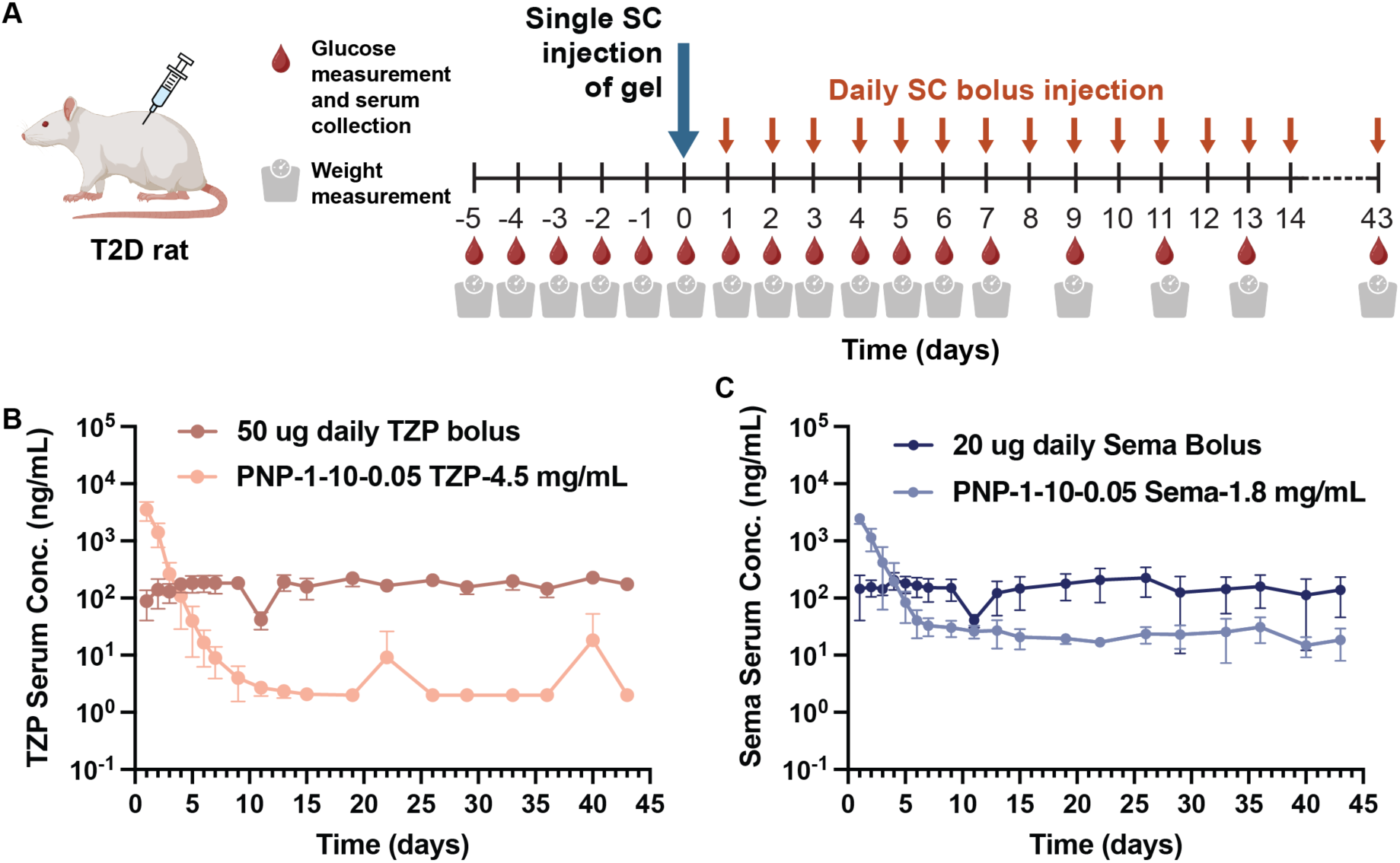
In vivo evaluation of tirzepatide and semaglutide release kinetics from PNP hydrogel formulations in diabetic rats. A single subcutaneous (SC) administration of TZP- or Sema-loaded PNP hydrogel sustained therapeutic drug levels for 6 weeks, in contrast to the short-acting effects of daily TZP (50 *μ*g) or Sema (20 *μ*g) injections. **(A)** Treatment timeline indicating dosing schedule, blood glucose (BG) monitoring, serum collection, and body weight measurements. Diabetic rats received either a single 500 *μ*L s.c. injection of PNP-1-10-0.05 hydrogel loaded with TZP (4.5 mg/mL) or Sema (1.8 mg/mL), or daily s.c. bolus injections of PBS, Sema (20 *μ*g/day), or TZP (50 *μ*g/day). **(B)** Pharmacokinetic profile of fasted male diabetic rats (5 ≤ n ≤ 6) treated with either daily TZP injections or a single TZP-loaded hydrogel injection. **(C)** Pharmacokinetic profile of male type 2 diabetic rats (5 ≤ n ≤ 6) treated with either daily Sema injections or a single Sema-loaded hydrogel injection.

Following the onset of diabetes,^[45, 46]^ rats were fasted overnight to ensure an average starting blood glucose (BG) level of approximately 130 mg dL^−1^ to 400 mg dL^−1^.^[47]^ Fasting BG levels were measured once a day for five days before treatment and rats were considered diabetic when at least three out of five fasting BG measurements, taken over the course of five days, fell between 130–400 mg dL^−1^ (**Figure 4A**). We validated previously reported pharmacokinetic parameters^[48]^ in diabetic rats (i.e., elimination half-life, volume of distribution, bioavailability, and absorption rate) by conducting a 24-hour pharmacokinetic (PK) study following intravenous (IV) administration of a 20 *µ*g dose of Sema in a saline vehicle (**Table S1**). We conducted an oral glucose tolerance test (OGTT) to ensure similar glucose response when grouping the rats into treatment groups (**Method S3**). Based on the OGTT, rats with similar glucose tolerance profile were paired, then these rats were randomized into treatment groups (**Figure S2**). We then compared the PK following subcutaneous administration of four treatment regimens: (i) repeated daily 20 *µ*g injections of Sema in a saline vehicle to mimic current clinical practice,^[48]^ (ii) single PNP Sema hydrogel formulations, (iii) single PNP TZP hydrogel formulations, and (iv) repeated daily PBS injections as an untreated control (5 ≤ n ≤ 6 for each treatment group). Sema has a half-life of 7 days in humans at a typical dose of 0.5 to 1 mg,^[49]^ and PK modeling shows that daily 20 *µ*g dosing in rats recapitulates the current clinical treatment regimen for patients.^[23]^ In contrast, the single hydrogel treatment groups represent our approach to long-acting GLP-1 RA delivery. For the first seven days after the start of treatment, and three times a week thereafter, serum was collected for PK analysis and BG was measured (**Figure 4A**).

The serum concentrations of TZP/Sema were measured over time following the subcutaneous administration of each treatment by ELISA to assess the pharmacokinetic profiles of each hydrogel-based formulation. We hypothesized that our hydrogel formulations would maintain therapeutically relevant concentrations of TZP/Sema, comparable to daily 20 *µ*g TZP/Sema administration, for six weeks in this rat model. Indeed, all hydrogel formulations we evaluated in this study effectively maintained relevant concentrations of TZP/Sema throughout the duration of the six-week-long study (**Figure 4B, C**). In **Figure 4D**, the HbA1c data indicate that both PNP hydrogel formulations of TZP (4.5 mg/mL) and Sema (1.8 mg/mL) effectively maintained or modestly reduced glycemic levels over 6 weeks following a single injection, demonstrating sustained therapeutic activity. In contrast, daily bolus injections of TZP/Sema showed greater inter-animal variability, suggesting less consistent glycemic control.

One important metric for adverse effects of GLP-1 RA therapies (typically gastrointestinal discomfort when initiating treatment) is Cmax, which is the maximum concentration of GLP-1 RA in the serum following treatment. In our studies, daily 50 *μ*g TZP administration yielded 162 ± 36 ng/mL over 45 days, while daily 20 *μ*g Sema administration yielded an average serum concentration of 152 ± 51 ng/mL during the same duration. Serum levels were relatively stable over the course of the treatment period with no clear peaks in the PK profile, which is typical of existing clinical administration regimens. In contrast, both TZP and Sema-loaded hydrogels reached Cmax values within 1 day of treatment, followed by a continuous, slow decrease in serum concentrations over the course of the first week to a steady state that remained throughout the duration of the study. The Cmax values for the gel treatment groups were 2200 ± 290 ng/mL (PNP-1-10-0.05-TZP) (**Figure 4B**, **Table S2**) and 3500 ± 1200 ng/mL (PNP-1-10-0.05-Sema) (**Figure 4B**, **Table S2**). While the Cmax values observed on the first day for all hydrogel-based TZP and Sema formulations might be expected to cause nausea and suppressed food intake in rodents, no changes to animal behavior were observed throughout the study.

We also examined the ability of the hydrogel-based long-acting TZP formulations to reduce average BG after each 6-week treatment regimen. In this study, all GLP-1 RA treatments resulted in a significant reduction in average BG over the course of the study, whereas untreated PBS controls experienced no significant change in average BG (**Figure 5A–B**). The daily dosing of 20 *μ*g Sema resulted in a 14 ± 4% reduction in average BG over the course of the 6-week study.

**Figure 5.**
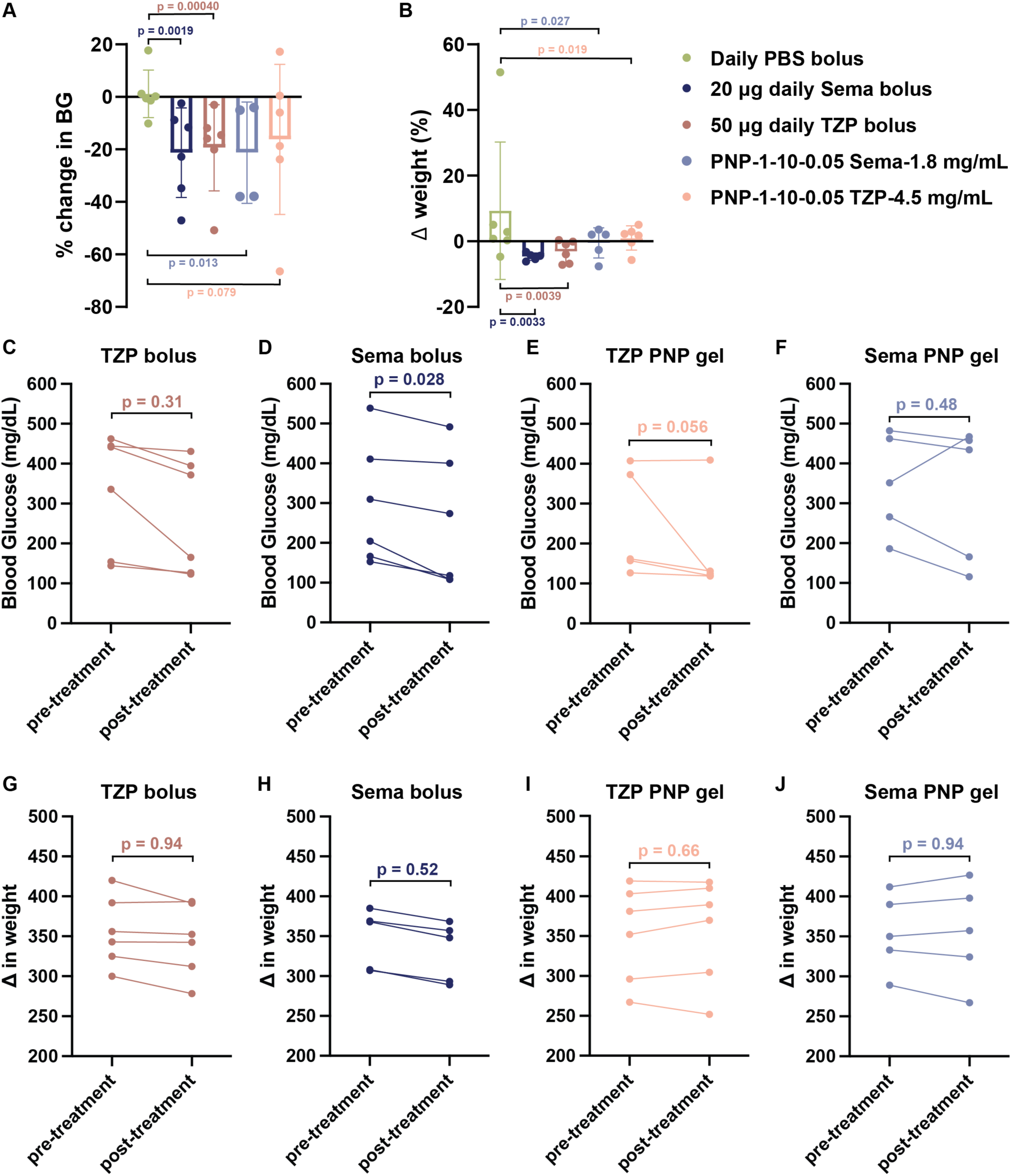
Therapeutic efficacy of a single PNP hydrogel injection in lowering blood glucose and limiting weight gain in diabetic rats. **(A)** Blood glucose (BG) levels over 6 weeks in type 2-like diabetic male rats (5 ≤ n ≤ 6) following treatment with a single subcutaneous administration of drug-loaded PNP hydrogel or daily bolus injections of Sema (20 *μ*g/day), TZP (50 *μ*g/day), or PBS. All hydrogel and daily treatment groups showed significantly reduced BG compared to the control group that receives daily PBS bolus SC injection (*p < 0.05). **(B)** Body weight changes over 6 weeks indicate that a single hydrogel administration results in similar or improved weight control compared to daily Sema injections. **(C–F)** Individual BG traces for each rat pre- and post-treatment across treatment groups. **(G–J)** Individual body weight traces for each rat pre- and post-treatment across treatment groups. (5 ≤ n ≤ 6)

**Figures 5C–F** and **5G–J** illustrate the pharmacodynamic impact of Sema and TZP delivered via PNP hydrogels, compared to their respective bolus injections. In **Figures 5C** and **D**, Sema bolus and PNP-1-10-0.05 Sema-1.8 mg/mL hydrogel both induced reductions in blood glucose, but the hydrogel formulation (**Figure 5D**) exhibited more durable glycemic control over time, with lower inter-animal variability. **Figures 5E** and **F** present corresponding body weight trends, showing that PNP-1-10-0.05 Sema-1.8 mg/mL mitigated the acute weight loss observed with bolus Sema, supporting a more stable metabolic profile. Similarly, **Figures 5G** and **H** show that PNP-1-10-0.05 Sema-1.8 mg/mL hydrogel resulted in sustained glucose lowering compared to the bolus injection, maintaining efficacy across the cohort. Body weight trajectories (**Figure 5I, J**) reveal that while daily TZP bolus caused noticeable weight loss in some animals, the hydrogel formulation promoted a more gradual and controlled weight reduction, consistent with improved tolerability and depot stability. Together, these changes underscore the therapeutic advantage of hydrogel-based delivery, with PNP formulations enhancing both the consistency and duration of GLP-1 receptor agonist action.

Both TZP and Sema demonstrated effective glycemic control in diabetic rats when delivered via PNP hydrogel formulations, with notable differences in consistency and magnitude of therapeutic response. Bolus administration of either peptide led to reductions in blood glucose, though the effect of Sema was more variable across individuals. In contrast, TZP produced a more consistent and moderate reduction, aligning with its known clinical potency. Notably, sustained delivery via PNP hydrogels enhanced therapeutic efficacy and uniformity for both drugs. The PNP 1-10-0.05 TZP-4.5 mg/mL formulation yielded an average reduction in blood glucose of approximately 40 mg/dL, indicating sustained bioactivity and controlled release. The PNP 1-10-0.05 Sema-1.8 mg/mL formulation similarly lowered glucose levels with an average reduction of about 68 mg/dL, though with greater inter-animal variability. (**Figure 5A**) Body weight outcomes further differentiated the delivery strategies. While bolus administration, especially with TZP, was associated with acute weight loss, hydrogel-based formulations mitigated this effect, supporting more stable weight profiles throughout the treatment period (**Figure 5B**). These findings suggest that PNP hydrogels provide a tunable and biocompatible platform for the sustained release of GLP-1 receptor agonists. TZP demonstrated more consistent pharmacodynamic effects than Sema in hydrogel form, reinforcing its promise as a candidate for long-acting injectable therapies in the management of type 2 diabetes.

An oral glucose tolerance test (OGTT) was performed prior to treatment and 43 days after the treatment to assess baseline glycemic control and ensure comparable glucose responsiveness across all treatment groups. This stratification is essential for minimizing variability due to differences in diabetes severity, enabling more accurate comparisons of treatment efficacy. As shown in the pre-treatment AUC data (**Figure S4A**, **C**, **Method S3**), all groups demonstrated elevated blood glucose excursions, consistent with impaired glucose tolerance. Notably, after 43 days of treatment, significant reductions in AUC were observed in all groups receiving GLP-1 receptor agonists, indicating improved glycemic control (**Figure S4B**, **D**, **Method S3**). The group receiving daily 20 *μ*g Sema showed a modest reduction in AUC, while the hydrogel-treated PNP-1-10-0.05 Sema-1.8 mg/mL group achieved comparable or even superior glucose tolerance, suggesting effective sustained release. The PNP-1-10-0.05 TZP-4.5 mg/mL hydrogel group demonstrated the largest drop in AUC, reflecting the potent and prolonged glucose-lowering effect of TZP when delivered via hydrogel. In contrast, the PBS-treated group showed minimal changes, reinforcing the therapeutic effect of both GLP-1 analogs. Collectively, these results validate the utility of GTT as a functional readout of treatment efficacy and demonstrate the hydrogel formulations’ ability to restore glucose handling over an extended period.

### 2.5. Blood chemistry and pathological impact of GLP-1 RA PNP hydrogel therapeutics

To assess the systemic impact and biosafety of GLP-1 receptor agonist (GLP-1 RA) therapeutics delivered via PNP hydrogels compared to traditional bolus administration, comprehensive blood chemistry panels were collected from diabetic rats at baseline (Day -1) and at the conclusion of the study (Day 43) (**Figure 6**). In addition, necropsy was performed on Day 43 to evaluate biocompatibility and potential local or systemic toxicity. Tissues collected for histological analysis included the kidney (**Figure S3**), liver (**Figure S4**), pancreas (**Figure S5**), and skin at or near the injection site (**Figure S6**). As shown in Figure 6, the blood chemistry panels present pre- and post-treatment values for key clinical biomarkers collected at Day 0 prior to the injection and at Day 43 after the treatment, including hemoglobin A1c (HbA1c) (**Figure 6B**), aspartate aminotransferase (AST) (**Figure 6C**), alanine aminotransferase (ALT) (**Figure 6D**), total bilirubin (**Figure 6E**), creatinine (**Figure 6F**), and blood urea nitrogen (BUN) (**Figure 6G**). These markers enable evaluation of hepatic and renal function, systemic inflammation, and long-term glycemic controls.

**Figure 6.**
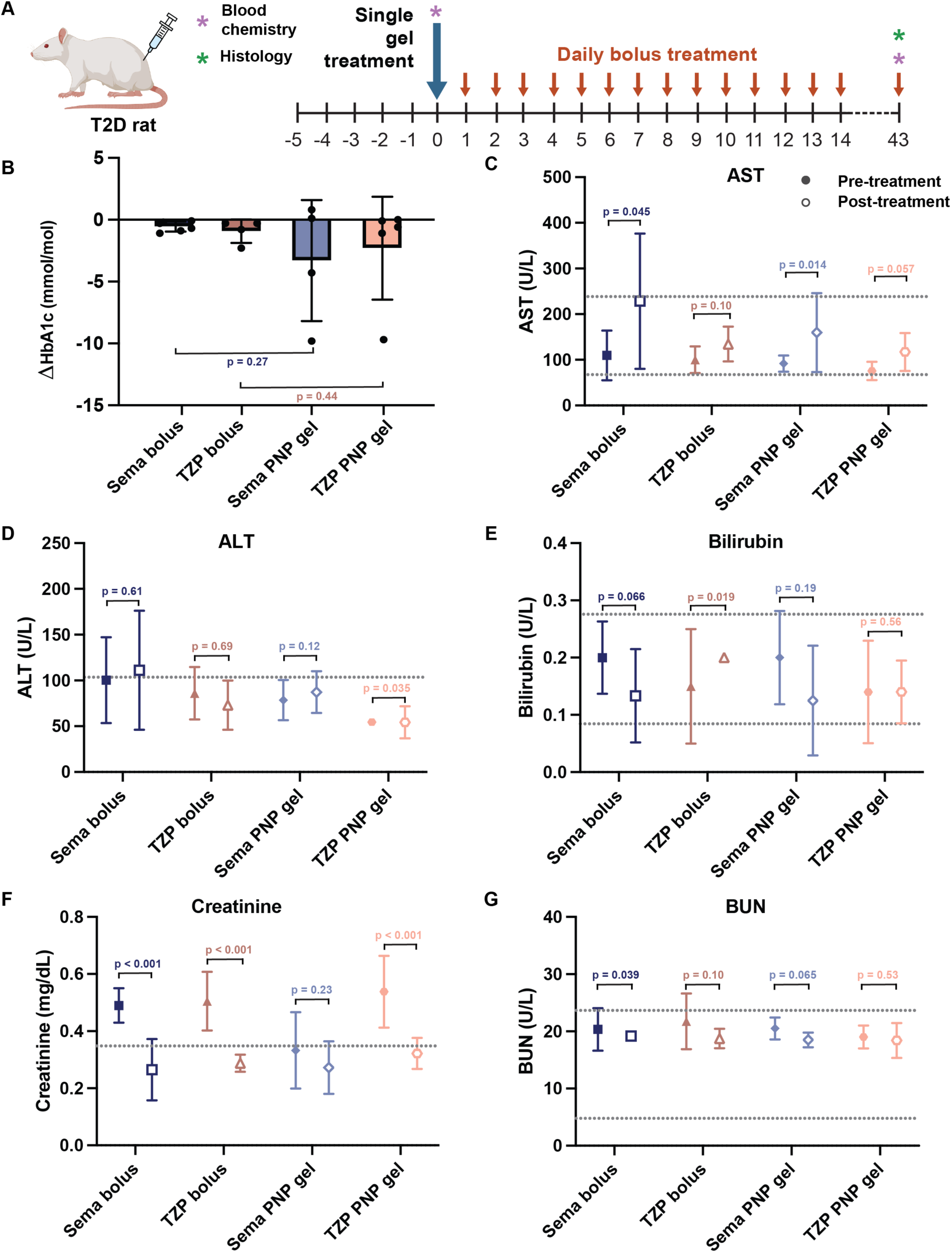
Blood chemistry profiles in T2D rats pre- and post-treatment with daily bolus or one injection of PNP hydrogel formulation of semaglutide or tirzepatide. **(A)** Blood chemistry parameters were assessed before and after treatment (Day 0 prior to the injection vs. Day 43) in type 2 diabetic rats receiving either daily PBS, daily Sema injections at 20 *μ*g or 50 *μ*g, a single 500 *μ*L injection of PNP-1-10-0.05 Sema-1.8 mg/mL, or a single injection of PNP-1-10-0.05 TZP, 4.5 mg/mL. Tissues were collected at the end of Day 43 for histology analysis. **(B)** Change in hemoglobin A1c (HbA1c) levels in T2D rats from pre-treatment to post-treatment, serving as a marker of glycemic control. The chemistry panels compare values from pre-treatment to Day 43 post-treatment for **(C)** aspartate aminotransferase (AST), **(D)** alanine aminotransferase (ALT), **(E)** total bilirubin, **(F)** creatinine, and **(G)** blood urea nitrogen (BUN). The dotted lines in each plot indicate the range of values exhibited in a healthy population of Sprague Dawley rats (age 8-9 weeks), defined as the mean ± two standard deviations.^[52]^ Error bars, mean ± s.e. (5 ≤ n ≤ 6)

**Figures 6C** and **D** present serum levels of AST and ALT, two standard biomarkers used to evaluate hepatic function and detect hepatocellular injury.^[50]^ While no treatment group exhibited ALT elevations beyond clinically concerning threshold, all groups showed slight increase in ALT following administration. Notably, the Sema treatment groups demonstrated statistically significant elevations in ALT, whereas the TZP groups did not (**Figure 6C**, **D**). Prolonged subcutaneous residence of GLP-1 RAs within the PNP hydrogel matrix did not induce overt hepatotoxicity and suggest that TZP may be more favorable in maintaining hepatic stability in this hydrogel-based delivery platform.

**Figures 6B** and **E–G** provide longitudinal analyses of additional metabolic and clinical safety markers, including HbA1c, total bilirubin, creatinine, and BUN, to assess systemic effects and therapeutic efficacy.^[51]^ Both hydrogel-treated groups maintained bilirubin and BUN levels comparable to or below baseline, with no indication of liver or kidney dysfunction throughout the 6-week study period. Serum creatinine levels remained stable or slightly declined in animals receiving PNP-based formulations, whereas bolus-treated groups showed greater variability. This suggests that PNP hydrogels may confer improved renal safety profiles. Notably, the PNP-1-10-0.05 TZP hydrogel group exhibited the most stable HbA1c profiles across animals, highlighting its potential for predictable, long-term glucose regulation. While the Sema hydrogel formulation also reduced HbA1c levels in multiple animals, it showed slightly greater variability. Still, both PNP-based depots of Sema at 1.8 mg/mL and TZP at 4.5 mg/mL maintained or modestly lowered HbA1c, consistent with sustained therapeutic exposure and metabolic control. The similarity in glycemic outcomes between daily injection and single-dose hydrogel groups underscores the comparable efficacy of both delivery methods, with the hydrogel strategy offering a distinct advantage in dosing convenience to reduce administration burdens and improve patient compliance.

These results collectively affirm the safety, efficacy, and biocompatibility of PNP hydrogels as a long-acting injectable platform for GLP-1 RAs. Importantly, the ability to deliver both Sema and TZP, two structurally and mechanistically distinct incretin mimetics, demonstrates the versatility and translatability of the PNP hydrogel system across therapeutic peptides. This platform holds promise for simplifying chronic diabetes management through reduced dosing frequency, consistent pharmacodynamic effects, and broad applicability to next-generation peptide therapeutics.

Histological evaluation of liver, pancreas, kidney, and skin tissues at Day 43 post-treatment revealed improved tissue compatibility with PNP-tween hydrogel-based delivery systems relative to daily bolus injections. Among all formulations, PNP-1-10-0.05 Sema (1.8 mg/mL) consistently preserved normal tissue architecture with minimal signs of inflammation or cellular stress. In the liver, the tissues maintained intact hepatic structure with limited vacuolization or immune infiltration. Conversely, daily TZP bolus induced mild hepatocellular vacuolation and perivascular inflammation, which was partially mitigated in the PNP-1-10-0.05 TZP (4.5 mg/mL) formulation group.

Pancreatic tissue from the PNP-1-10-0.05 Sema gel group exhibited well-preserved islet morphology and minimal acinar disruption, in contrast to daily TZP bolus treatment group, which showed moderate exocrine stress and focal necrosis. Similarly, kidney and dermal tissues from the PNP gel-treated animals displayed reduced tubular degeneration and dermal remodeling. Though PNP-1-10-0.05 TZP exhibited slightly greater tissue response than the PNP-1-10-0.05 Sema gel, the TZP gel group still offered improved histological profiles relative to bolus injection. Overall, these findings suggest that PNP-based depots mitigate both systemic and local tissue stress, offering a biocompatible platform for sustained GLP-1 receptor agonist delivery and presenting a promising strategy for long-acting TZP administration that balances efficacy and safety.

## 3. Conclusion

This study demonstrates the adaptability and translational potential of PNP hydrogel systems for the sustained subcutaneous delivery of GLP-1 receptor agonists, with a specific focus on TZP and Sema. By leveraging the modularity of the PNP hydrogel platform, we developed depot formulations capable of delivering clinically relevant doses of both incretin-based therapeutics over an extended duration following a single injection. Rheological characterization confirmed that incorporation of either TZP or Sema did not significantly alter the mechanical properties of the hydrogel, supporting formulation flexibility and robustness. In vitro release assays and in vivo pharmacokinetics in a diabetic rat model revealed ultra-sustained drug release with stable therapeutic plasma concentrations over six weeks, mirroring or surpassing the efficacy of conventional daily injections.

Both PNP-1-10-0.05 Sema-1.8 mg/mL and PNP-1-10-0.05 TZP-4.5 mg/mL formulations yielded durable glycemic control and stabilized body weight, confirming the pharmacodynamic effectiveness of the depot platform. Notably, TZP exhibited more consistent pharmacokinetic and glycemic response profiles, reinforcing its promise as a candidate for long-acting injectable therapies. Blood chemistry panels and histological analyses of liver, pancreas, kidney, and skin revealed that PNP-based delivery was well tolerated, with reduced organ-level inflammation and tissue remodeling relative to daily bolus administration. Among the tested formulations, PNP-1-10-0.05 Sema-1.8 mg/mL exhibited the most favorable safety profile across all evaluated tissues, while PNP-1-10-0.05 TZP-4.5 mg/mL, though eliciting a modest increase in localized tissue response, remained substantially more biocompatible than daily bolus TZP.

Altogether, these findings underscore the clinical promise of PNP hydrogels as an injectable, scalable, and tunable platform for the ultra-sustained delivery of incretin-based therapeutics. This platform has the potential to improve patient adherence, reduce dosing frequency, and mitigate off-target toxicity in chronic disease management. Future work will explore further optimization of drug release kinetics, long-term safety across larger animal models, and eventual clinical translation of this technology for type 2 diabetes and metabolic disorders.

## 4. Materials and Methods

### Materials

Tween-20 (polyethylene glycol sorbitan monolaurate) was obtained from Sigma-Aldrich and used without further purification. Hydroxypropyl methylcellulose (HPMC, USP grade), N,N-diisopropylethylamine (Hunig’s base), hexanes, diethyl ether, N-methyl-2-pyrrolidone (NMP), dichloromethane (DCM), lactide (LA), 1-dodecyl isocyanate, and 1,8-diazabicyclo[5.4.0]undec-7-ene (DBU) were also purchased from Sigma-Aldrich and used as received. Monomethoxy polyethylene glycol (PEG, 5 kDa) was obtained from Sigma-Aldrich and dried under vacuum prior to use. All glassware and stir bars were oven-dried at 180 °C. Where indicated, solvents were degassed using three freeze–pump–thaw cycles. Pharmaceutical-grade semaglutide (Ozempic^®^) and tirzepatide (Mounjaro^®^) were procured from Stanford University’s pharmacy. To remove formulation excipients, the drug solutions were dialyzed against deionized water for 5 days and subsequently lyophilized.

### Preparations of HPMC-C12

Dodecyl-modified hydroxypropyl methylcellulose (HPMC-C12) was synthesized following previously reported protocols.^[53]^ Briefly, HPMC (1.0 g) was dissolved in 40 mL of N-methyl-2-pyrrolidone (NMP) under constant stirring at 80 °C for 1 h. After cooling the solution to room temperature (RT), a separate solution containing 1-dodecylisocyanate (105 mg, 0.5 mmol) and N,N-diisopropylethylamine (Hunig’s base, 3 drops) in 5 mL of NMP was prepared and added dropwise to the polymer solution. The reaction mixture was stirred at RT for 16 h to allow complete functionalization. The resulting product was precipitated into acetone, decanted, redissolved in deionized water (2 wt%), and dialyzed against water for 3 ∼ 4 days using a dialysis membrane (MWCO: 12–14 kDa) to remove residual reagents and solvent. The dialyzed polymer was subsequently lyophilized and reconstituted in sterile phosphate-buffered saline (PBS) to a final concentration of 60 mg mL⁻¹ for use in hydrogel formulations.

### Preparation of PEG-PLA NPs

PEG-PLA was prepared as previously reported.^[53]^ Monomethoxy-PEG (5 kDa, 0.25 g, 4.1 mmol) and DBU (15 *µ*L, 0.1 mmol; 1.4 mol% relative to LA) were dissolved in anhydrous dichloromethane (1.0 mL). LA (1.0 g, 6.9 mmol) was dissolved in anhydrous DCM (3.0 mL) with mild heating. The LA solution was added rapidly to the PEG/DBU solution and was allowed to stir for 8 min. The reaction mixture was quenched and precipitated by a 1:1 hexane and ethyl ether solution. The synthesized PEG−PLA was collected and dried under vacuum. Hydrogel permeation chromatography (GPC) was used to verify that the molecular weight and dispersity of polymers meet our quality control (QC) parameters. NPs were prepared as previously reported.^[53]^ A 1 mL solution of PEG-PLA in acetonitrile (50 mg mL^−1^) was added dropwise to 10 mL of water at RT under a high stir rate (600 rpm). NPs were purified by centrifugation over a filter (molecular weight cutoff of 10 kDa; Millipore Amicon Ultra-15) followed by resuspension in PBS to a final concentration of 200 mg mL^−1^. NPs were characterized by dynamic light scattering (DLS) to find the NP diameter, 37 ± 4 nm.

### PNP hydrogel preparation

Hydrogel formulations contained either 2 wt% HPMC-C12 and 10 wt% PEG-PLA NPs in PBS, or 1 wt% HPMC-C12 and 10 wt% PEG−PLA NPs in PBS. These hydrogels were made by mixing a 2:3:1 weight ratio of 6 wt% HPMC-C12 polymer solution, 20 wt% NP solution, and PBS containing GLP-1 RAs. The NP and aqueous components were loaded into one syringe, the HPMC-C12 was loaded into a second syringe and components were mixed using an elbow connector. After mixing, the elbow was replaced with a 21-gauge needle for injection.

### Injection force measurements

Injection force was assessed using a syringe pump-driven protocol adapted from previously established methods in our previous works.^[21, 43]^ A 1 mL syringe (BD) equipped with a 40 mm, 21-gauge needle was mounted onto a syringe pump configured to deliver fluid at a controlled volumetric flow rate based on the barrel diameter. A calibrated load cell (FUTEK LLB300) was attached to the syringe assembly to continuously measure the axial force required for extrusion. Force data were recorded in real time using a custom LabView interface. Injections proceeded until the measured force plateaued, at which point the pump was manually stopped.

### Rheological characterization of hydrogels

Rheological testing was performed at 25 °C using a 20-mm-diameter serrated parallel plate at a 600 *μ*m gap on a stress-controlled TA Instruments DHR-2 rheometer. All experiments were performed at 25 °C. Frequency sweeps were performed from 0.1 to 100 rad s^−1^ with a constant oscillation strain within the linear viscoelastic regime (1%). Amplitude sweeps were performed at a constant angular frequency of 10 rad s^−1^ from 0.01% to 10000% strain with a gap height of 500 *µ*m. Flow sweeps were performed from low to high stress with steady-state sensing. Steady shear experiments were performed by alternating between a low shear rate (0.1 s^−1^) and high shear rate (10 s^−1^) for 60 s each for three full cycles. Shear rate sweep experiments were performed from 10 to 0.001 s^−1^. Stress controlled yield stress measurements (stress sweeps) were performed from low to high stress with steady-state sensing and 10 points per decade.

### In vitro release

To evaluate release kinetics, 100 *μ*L of each hydrogel formulation was injected into four-inch glass capillary tubes. A 400 *μ*L volume of phosphate-buffered saline (PBS) was gently layered on top of the hydrogel to serve as the release medium. At designated time points (1, 3, 6, 12, 24, and 48 hours, and at 1 and 2 weeks), the PBS overlay was carefully collected for analysis and immediately replaced with fresh PBS to maintain infinite sink conditions. The concentration of semaglutide in each aliquot was quantified using a commercial enzyme-linked immunosorbent assay (ELISA, BMA BIOMEDICALS S-1530), enabling the construction of cumulative release profiles over time. (n = 3)

### Animal Studies

Animal studies were performed with the approval of the Stanford Administrative Panel on Laboratory Animal Care (APLAC-32873) and were in accordance with National Institutes of Health guidelines.

### In-vivo pharmacokinetics and pharmacodynamics in diabetic rats

For animals receiving long-acting therapies, a single 500 *μ*L subcutaneous injection of the PNP hydrogel formulation was administered on Day 0. In contrast, animals in the bolus treatment groups received daily 100 *μ*L subcutaneous injections of PBS, TZP, or Sema. Baseline blood samples were collected from the tail vein on Day 0 prior to treatment, and blood glucose levels were monitored daily for 43 days using a handheld glucometer (Bayer Contour Next).^[54]^ For pharmacokinetic analysis, tail vein blood samples were collected daily during the first seven days and twice weekly thereafter to quantify plasma concentrations of semaglutide or tirzepatide using ELISA (BMA BIOMEDICALS S-1530). Total drug exposure (AUC) and bioavailability were calculated at the study endpoint. Prior to treatment (Day −1), an oral glucose tolerance test (OGTT) was performed to evaluate baseline glucose responsiveness and to stratify animals into treatment groups with comparable glycemic profiles.^[46]^

### Biocompatibility

At the end of the 6-week study, rats were euthanized via carbon dioxide inhalation, and tissues (kidney, liver, pancreas, and skin) were collected for histological analysis. The liver was sectioned transversely at the left lateral and right medial lobes, and the kidney was sectioned longitudinally. Skin samples were harvested from the injection site. All tissues were fixed and processed for histology. Hematoxylin and eosin (H&E) staining was performed on all samples by Histo-tec Laboratory. In addition, skin sections were stained with Masson’s trichrome (TC) stain to evaluate collagen deposition and dermal structure. Histological assessments were conducted by two blinded observers with established expertise in rodent pathology. Assessment criteria were adapted based on organ-specific pathology: kidney sections were evaluated for tubular dilation, glomerular injury, and interstitial inflammation; liver sections for hepatocellular degeneration, steatosis, and inflammatory cell presence; and pancreas sections for acinar atrophy, islet disruption, and immune infiltration (3 ≤ n ≤ 5).

### Statistical analysis

All results are expressed as a mean ± standard deviation unless specified otherwise. For in-vivo experiments, Mead’s Resource Equation was used to identify a sample size above that additional subjects will have little impact on power. Comparison between groups was conducted with the Kolmogorov-Smirnov test in MATLAB to obtain the significance levels. A difference corresponding to values of p < 0.05 was considered statistically significant.

## Supporting Information

Supporting Information is available from the Wiley Online Library or from the author.

## Acknowledgements

This work was supported by the National Institute of Diabetes and Digestive and Kidney Diseases (NIDDK) (R01DK119254). A.I.D. was supported by a Schmidt Science Fellows Award. L.T.N. received support from a Stanford Graduate Fellowship in Science and Engineering. In memoriam, we dedicate this work to L.T.N., whose unwavering commitment to advancing diabetes research, scientific rigor, and collaborative spirit left a lasting impact on this project and the broader scientific community. In addition, J.Y., C.K.J., O.M.S., N.E., and C.M.W. were supported by National Science Foundation Graduate Research Fellowships. S.K. gratefully acknowledges support from the 2023 Stanford SURF program and mentorship from Appel Lab. We also thank all members of the Appel Lab for their valuable discussions and inputs throughout this project. The authors are grateful to the Stanford Animal Diagnostic Lab and the Veterinary Service Center staff for their technical assistance.

## Author Contributions

A.I.D. and E.A.A. conceived the project idea. A.I.D. and C.D. designed the experiments and performed most of the experimental work. Additional authors contributed to specific experiments, data analysis, and manuscript preparation. All authors reviewed and approved the final manuscript.

## Data Availability

The data that support the findings of this study are available from the corresponding author upon reasonable request.

## References

1. Williams, R., et al., *Global and regional estimates and projections of diabetes-related health expenditure: Results from the International Diabetes Federation Diabetes Atlas*, *9th edition*. Diabetes Research and Clinical Practice, 2020. 162.

2. CDC, Centers for Disease Control and Prevention. 2022, National Diabetes Statistics Report website: https://www.cdc.gov/diabetes/data/statistics-report/index.html

3. DeFronzo, R.A., et al., Type 2 diabetes mellitus. Nature Reviews Disease Primers, 2015. 1(1): p. 15019.

4. White, M.G., J.A.M. Shaw, and R. Taylor, Type 2 Diabetes: The Pathologic Basis of Reversible β-Cell Dysfunction. Diabetes Care, 2016. 39(11): p. 2080–2088.

5. Mezza, T., et al., β-Cell Fate in Human Insulin Resistance and Type 2 Diabetes: A Perspective on Islet Plasticity. Diabetes, 2019. 68(6): p. 1121–1129.

6. De Block, C., et al., Tirzepatide for the treatment of adults with type 2 diabetes: An endocrine perspective. Diabetes Obes Metab, 2023. 25(1): p. 3–17.

7. Skyler, J.S., et al., Differentiation of Diabetes by Pathophysiology, Natural History, and Prognosis. Diabetes, 2017. 66(2): p. 241–255.

8. Chung, W.K., et al., Precision medicine in diabetes: a Consensus Report from the American Diabetes Association (ADA) and the European Association for the Study of Diabetes (EASD). Diabetologia, 2020. 63(9): p. 1671–1693.

9. Taskinen, M.R. and J. Borén, New insights into the pathophysiology of dyslipidemia in type 2 diabetes. Atherosclerosis, 2015. 239(2): p. 483–95.

10. Fonseca, V.A., Defining and Characterizing the Progression of Type 2 Diabetes. Diabetes Care, 2009. 32(suppl_2): p. S151–S156.

11. Moura, F.A., B.M. Scirica, and C.T. Ruff, Tirzepatide for diabetes: on track to SURPASS current therapy. Nature Medicine, 2022. 28(3): p. 450–451.

12. Nauck, M.A., et al., GLP-1 receptor agonists in the treatment of type 2 diabetes - state-of-the-art. Mol Metab, 2021. 46: p. 101102.

13. Christensen, M.B., et al., Glucose-dependent Insulinotropic Polypeptide: Blood Glucose Stabilizing Effects in Patients With Type 2 Diabetes. The Journal of Clinical Endocrinology & Metabolism, 2014. 99(3): p. E418–E426.

14. Nauck, M.A., et al., The evolving story of incretins (GIP and GLP-1) in metabolic and cardiovascular disease: A pathophysiological update. Diabetes, Obesity and Metabolism, 2021. 23(S3): p. 5–29.

15. DiMatteo, M.R., et al., Patient adherence and medical treatment outcomes: a meta-analysis. Med Care, 2002. 40(9): p. 794–811.

16. Pagès-Puigdemont, N., et al., Patients’ Perspective of Medication Adherence in Chronic Conditions: A Qualitative Study. Adv Ther, 2016. 33(10): p. 1740–1754.

17. Kim, J., et al., Medication adherence: The elephant in the room. U.S. Pharmacist, 2018. 43: p. 30–34.

18. Medication Adherence - Improving Health Outcomes. American College of Preventative Medicine, 2011. www.acpm.org/?MedAdherTT_ClinRef. Accessed April 9, 2024.

19. Lau, D.T. and D.P. Nau, Oral antihyperglycemic medication nonadherence and subsequent hospitalization among individuals with type 2 diabetes. Diabetes Care, 2004. 27(9): p. 2149–53.

20. Baryakova, T.H., et al., Overcoming barriers to patient adherence: the case for developing innovative drug delivery systems. Nature Reviews Drug Discovery, 2023. 22(5): p. 387–409.

21. Jons, C.K., et al., Yield-Stress and Creep Control Depot Formation and Persistence of Injectable Hydrogels Following Subcutaneous Administration. Advanced Functional Materials. n/a(n/a): p. 2203402.

22. Sen, S., et al., Evolving transport properties of dynamic hydrogels enable self-tuning of short-and long-term cargo delivery. arXiv preprint arXiv:2503.14915, 2025.

23. Appel, E.A., et al., Self-assembled hydrogels utilizing polymer–nanoparticle interactions. Nature Communications, 2015. 6(1): p. 6295.

24. Grosskopf, A.K., et al., Injectable supramolecular polymer–nanoparticle hydrogels enhance human mesenchymal stem cell delivery. Bioengineering & Translational Medicine, 2020. 5(1): p. e10147.

25. Stapleton, L.M., et al., Use of a supramolecular polymeric hydrogel as an effective post-operative pericardial adhesion barrier. Nature Biomedical Engineering, 2019. 3(8): p. 611–620.

26. Steele, A.N., et al., A Biocompatible Therapeutic Catheter-Deliverable Hydrogel for In Situ Tissue Engineering. Advanced Healthcare Materials, 2019. 8(5): p. 1801147.

27. Yu, A.C., et al., Physical networks from entropy-driven non-covalent interactions. Nature Communications, 2021. 12(1): p. 746.

28. Meis, C.M., et al., Injectable Supramolecular Polymer-Nanoparticle Hydrogels for Cell and Drug Delivery Applications. JoVE, 2021(168): p. e62234.

29. Liong, C.S., et al., Enhanced Humoral Immune Response by High Density TLR Agonist Presentation on Hyperbranched Polymers. Advanced Therapeutics, 2021. n/a(n/a): p. 2000081.

30. Stapleton, L.M., et al., Dynamic Hydrogels for Prevention of Post-Operative Peritoneal Adhesions. Advanced Therapeutics, 2021. n/a(n/a): p. 2000242.

31. Meany, E.L., et al., Generation of an inflammatory niche in an injectable hydrogel depot through recruitment of key immune cells improves efficacy of mRNA vaccines. bioRxiv, 2024: p. 2024.07. 05.602305.

32. Saouaf, O.M., et al., Sustained vaccine exposure elicits more rapid, consistent, and broad humoral immune responses to multivalent influenza vaccines. Advanced Science, 2024: p. 2404498.

33. d’Aquino, A.I., et al., Use of a biomimetic hydrogel depot technology for sustained delivery of GLP-1 receptor agonists reduces burden of diabetes management. Cell Rep Med, 2023. 4(11): p. 101292.

34. Kasse, C.M., et al., Subcutaneous delivery of an antibody against SARS-CoV-2 from a supramolecular hydrogel depot. bioRxiv, 2022: p. 2022.05.24.493347.

35. Grosskopf, A.K., et al., Delivery of CAR-T cells in a transient injectable stimulatory hydrogel niche improves treatment of solid tumors. Science Advances, 2022. 8(14): p. eabn8264.

36. Dong, C., et al., Water-Enhancing Gels Exhibiting Heat-Activated Formation of Silica Aerogels for Protection of Critical Infrastructure During Catastrophic Wildfire. Advanced Materials, 2024. 36(40): p. 2407375.

37. Changxin Dong, S.S., Zhennan Ru, Athena Kolli, Paxton S. Appel, Jonathan Fan, Eric A. Appel, Hydrogel-to-Aerogel Transitions in Polymer-Particle Hydrogels Expand the Wildfire Defense Window. arXiv:2503.14923, 2025.

38. d’Aquino, A.I., et al., Use of a biomimetic hydrogel depot technology for sustained delivery of GLP-1 receptor agonists reduces burden of diabetes management. Cell Reports Medicine, 2023. 4(11).

39. Frías, J.P., et al., Tirzepatide versus Semaglutide Once Weekly in Patients with Type 2 Diabetes. New England Journal of Medicine, 2021. 385(6): p. 503–515.

40. Ding, Y., et al., Evaluation and comparison of efficacy and safety of tirzepatide and semaglutide in patients with type 2 diabetes mellitus: A Bayesian network meta-analysis. Pharmacological Research, 2024. 199: p. 107031.

41. Wu, C., et al., Disruption of Escherichia coli amyloid-integrated biofilm formation at the air-liquid interface by a polysorbate surfactant. Langmuir, 2013. 29(3): p. 920–6.

42. Patapoff, T.W. and O. Esue, Polysorbate 20 prevents the precipitation of a monoclonal antibody during shear. Pharm Dev Technol, 2009. 14(6): p. 659–64.

43. Lopez Hernandez, H., J.W. Souza, and E.A. Appel, A Quantitative Description for Designing the Extrudability of Shear-Thinning Physical Hydrogels. Macromolecular Bioscience, 2021. 21(2): p. 2000295.

44. Szkudelski, T., Streptozotocin–nicotinamide-induced diabetes in the rat. Characteristics of the experimental model. Experimental Biology and Medicine, 2012. 237(5): p. 481–490.

45. Masiello, P., et al., Experimental NIDDM: development of a new model in adult rats administered streptozotocin and nicotinamide. Diabetes, 1998. 47(2): p. 224–9.

46. Chen, T., L. Kagan, and D.E. Mager, Population pharmacodynamic modeling of exenatide after 2-week treatment in STZ/NA diabetic rats. Journal of pharmaceutical sciences, 2013. 102(10): p. 3844–3851.

47. Wu, K.K. and Y. Huan, Streptozotocin-induced diabetic models in mice and rats. Curr Protoc Pharmacol, 2008. Chapter 5: p. Unit 5.47.

48. Knudsen, L.B. and J. Lau, The Discovery and Development of Liraglutide and Semaglutide. Frontiers in endocrinology, 2019. 10: p. 155–155.

49. Hall, S., D. Isaacs, and J.N. Clements, Pharmacokinetics and Clinical Implications of Semaglutide: A New Glucagon-Like Peptide (GLP)-1 Receptor Agonist. Clinical Pharmacokinetics, 2018. 57(12): p. 1529–1538.

50. Thakur, S., et al., Biomarkers of hepatic toxicity: an overview. Current Therapeutic Research, 2024: p. 100737.

51. Lala, V., M. Zubair, and D. Minter, Liver function tests. StatPearls, 2023.

52. Mann, J.L., et al., An ultrafast insulin formulation enabled by high-throughput screening of engineered polymeric excipients. Science translational medicine, 2020. 12(550): p. eaba6676.

53. Meis, C.M., et al., *S*elf-Assembled, Dilution-Responsive Hydrogels for Enhanced Thermal Stability of Insulin Biopharmaceuticals. ACS Biomaterials Science & Engineering, 2021. 7(9): p. 4221–4229.

54. Galderisi, A., E. Schlissel, and E. Cengiz, Keeping Up with the Diabetes Technology: 2016 Endocrine Society Guidelines of Insulin Pump Therapy and Continuous Glucose Monitor Management of Diabetes. Curr Diab Rep, 2017. 17(11): p. 111.

