## Supplementary material for "Long-acting hydrogel-based depot formulations of tirzepatide and semaglutide for the management of type 2 diabetes and weight": SI

**Method S1. Tirzepatide quantification and determining in vitro % cumulative release**

Quantification of tirzepatide was determined by high performance liquid chromatography–mass spectrometry (HPLC–MS) using a Sciex Exion LCAD equipped with a Thermo beta-basic-4 50 x 2.1 mm, 5 µm column. The mass spectrometry was achieved using a Sciex Qtrap 6500+ with multiple-reaction-monitoring. The ionization mode was turbo ion spray with positive ionization. Samples were analyzed at 40 °C at a flow rate of 0.7 mL min$^{-1}$. The mobile phase was 0.1% formic acid in water and methanol. To prepare samples for injection, a 20 µL aliquot of diluted sample was treated with 150 µL of 85% acetonitrile with perchloric acid containing the internal standard (terfenadine). The mixture was shaken on a shaker for 15 minutes and subsequently centrifuged at 3220 g for 15 minutes. A 140 µL aliquot of the supernatant was transferred for injection to the LC/MS/MS. Calibration standards and quality control samples were prepared by spiking the test compound into a blank and then processed with the unknown samples in the same batch. In vitro cumulative release was determined by taking the ratio of $\frac{M_t}{M_{inf}}$, where $M_t$ is the amount of drug released at time *t*, and $M_{inf}$ is the total amount of drug within the gel.

**Method S2. Streptozotocin/Nicotinamide (STZ/NA) induced model of type-2 diabetes in rats**

Male Sprague Dawley rats (160–230 g, 8–10 weeks old; Charles River) were used for all in vivo experiments. All animal procedures were conducted in accordance with institutional and national guidelines for the care and use of laboratory animals and were approved by the Stanford Institutional Animal Care and Use Committee (IACUC Protocol #32873). To induce type 2 diabetes (T2D), rats were fasted for 6–8 hours prior to treatment and then administered nicotinamide (NA) and streptozotocin (STZ). NA was dissolved in sterile 1× PBS and delivered via intraperitoneal injection at a dose of 110 mg/kg. STZ was freshly prepared in sodium citrate buffer (10 mg/mL) and injected intraperitoneally at a dose of 65 mg kg$^{-1}$. Following STZ administration, rats received 10% sucrose in their drinking water for 24 hours to prevent acute hypoglycemia. Blood glucose (BG) levels were subsequently monitored daily via tail vein sampling using a handheld Bayer Contour Next glucometer. Rats were considered diabetic when they exhibited three or more consecutive non-fasting BG readings within the range of 130–400 mg dL$^{-1}$. The induction success rate was roughly 50%.

**Method S3. Oral glucose tolerance test (OGTT)**

An oral glucose tolerance test (OGTT) was conducted to assess baseline glucose handling prior to treatment. Rats were fasted for 12 hours before the procedure. Baseline blood glucose levels were recorded at -30, -10, and -5 minutes prior to glucose administration. A 20 wt% D-glucose solution was prepared and administered orally at a dosage of 2 g $kg^{-1}$ body weight using a polyethylene gastric gavage tube. Post-administration blood glucose levels were measured at 5, 15, 30, 45, 60, and 120 minutes using a handheld glucometer (Bayer Contour Next). Based on individual OGTT response profiles, rats were stratified into bins with comparable glucose tolerance and subsequently randomized into treatment groups to ensure balanced baseline glycemic control.

**Method S4. *In vivo* Pharmacokinetics modeling for IV and SC data**

A 24-hour IV PK study was conducted to validate the PK parameters of Semaglutide and Tirzapetide in rats, and a single-compartment model was fit to obtain the elimination half-life (clearance rate) of the drugs. The differential equation governing the PK is

$$\frac{d}{dt}M(t) = -k_{\text{elim}}M(t)$$

The IV PK data is fit to this equation. Here, $k_{\text{elim}}$ is the elimination rate constant, from which the elimination half-life is determined as $\tau_{\text{elim}} = \frac{\ln 2}{k_{\text{elim}}}$ . The SC gel data was also fit to the one-compartment model outlined above to obtain the half-lives of drug elimination from the body.

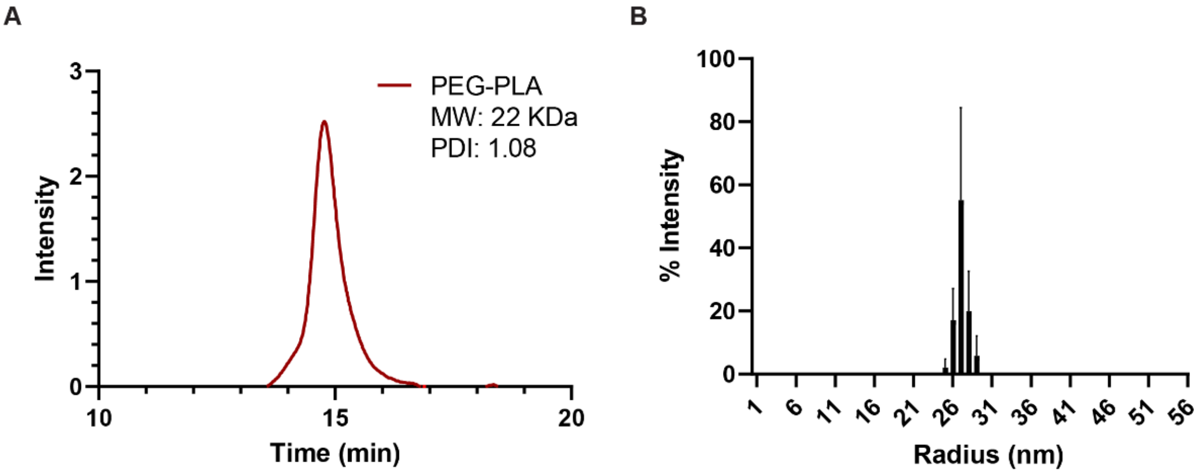

**Figure S1. Characterization of PEG-PLA polymer and nanoparticles. (A)** Size exclusion chromatography (SEC) trace of PEG-PLA block copolymer, displaying a single, monomodal peak with a number-average molecular weight (Mn) of 22 kDa and a dispersity (Đ) of 1.08. **(B)** Dynamic light scattering (DLS) analysis of PEG-PLA nanoparticles following nanoprecipitation, showing a hydrodynamic diameter ($D_h$) of 33.2 nm and a low polydispersity index (PDI = 0.038), indicating uniform particle size distribution.

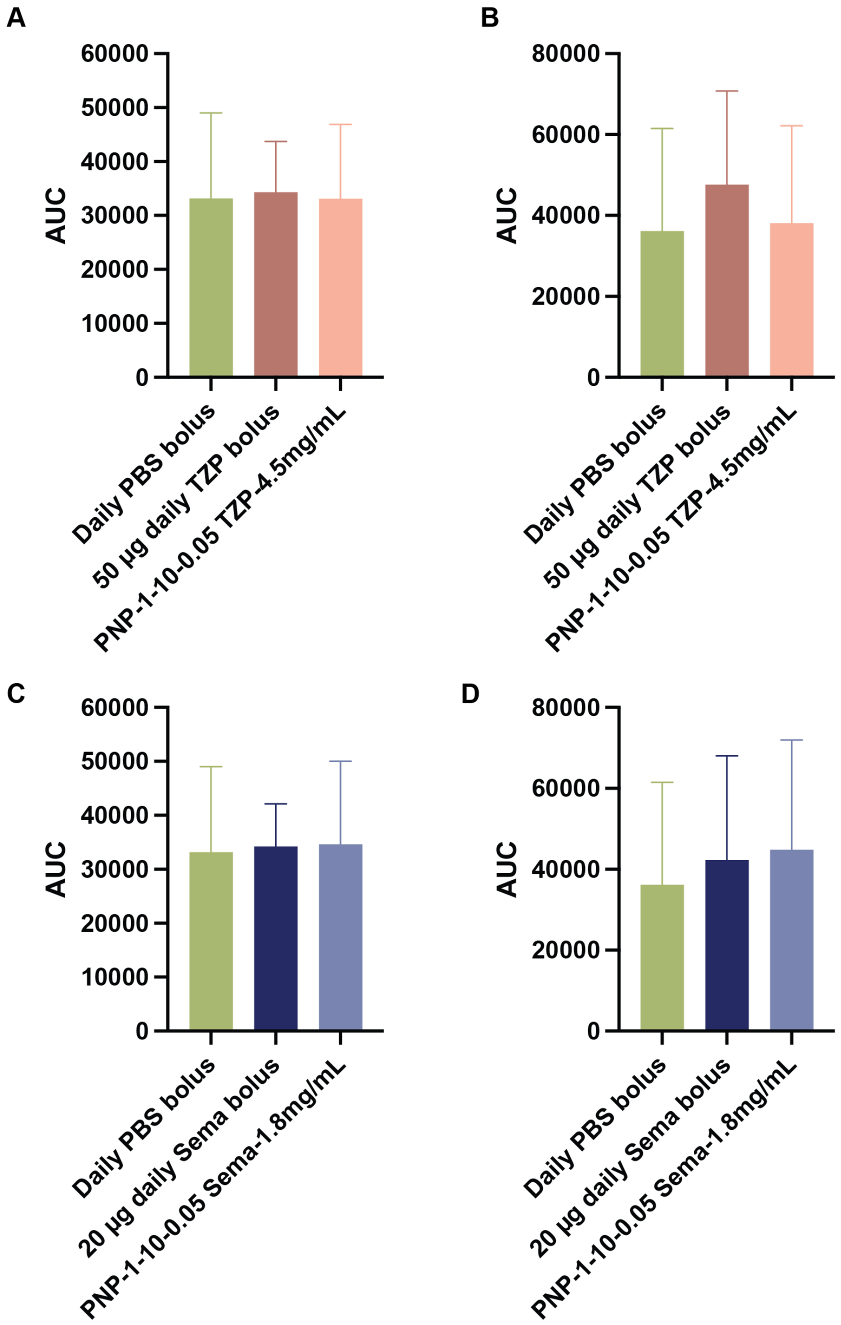

**Figure S2. Oral Glucose Tolerance Test (OGTT) for pre- and post-treatment comparison.** An OGTT was performed to assess glucose tolerance prior to treatment and to stratify rats into three treatment groups. Blood glucose levels were measured at baseline (−30, −10, and −5 min) and at 5, 15, 30, 45, 60, and 120 minutes following oral glucose administration (2 g $kg^{-1}$ body weight). Rats with comparable glucose tolerance, as determined by area under the curve (AUC) analysis, were paired and then randomized into treatment groups. **(A)** Pre-treatment OGTT results. **(B)** Post-treatment OGTT results.

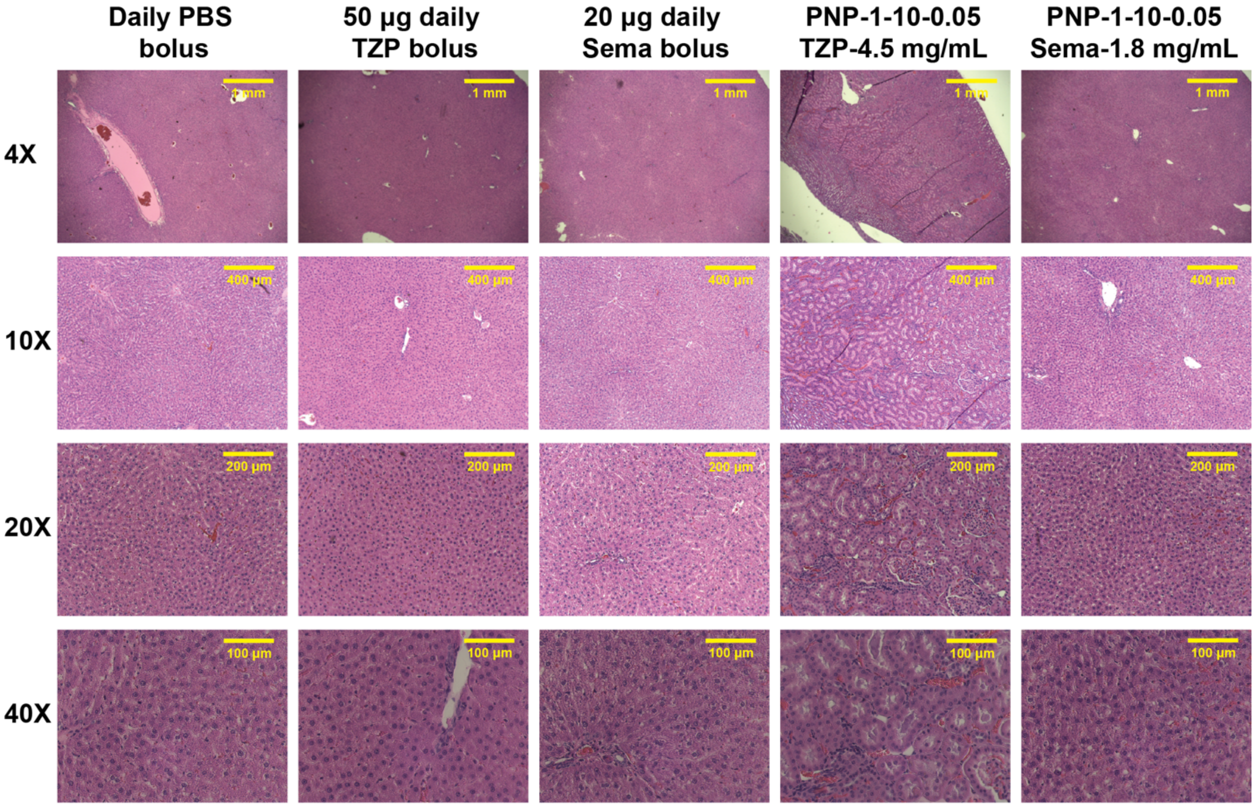

**Figure S3. Kidney Histology – Day 43.** Representative hematoxylin and eosin (H&E) stained cross-sections of kidney tissue collected on Day 43 post-treatment. Images illustrate kidney morphology across treatment groups.

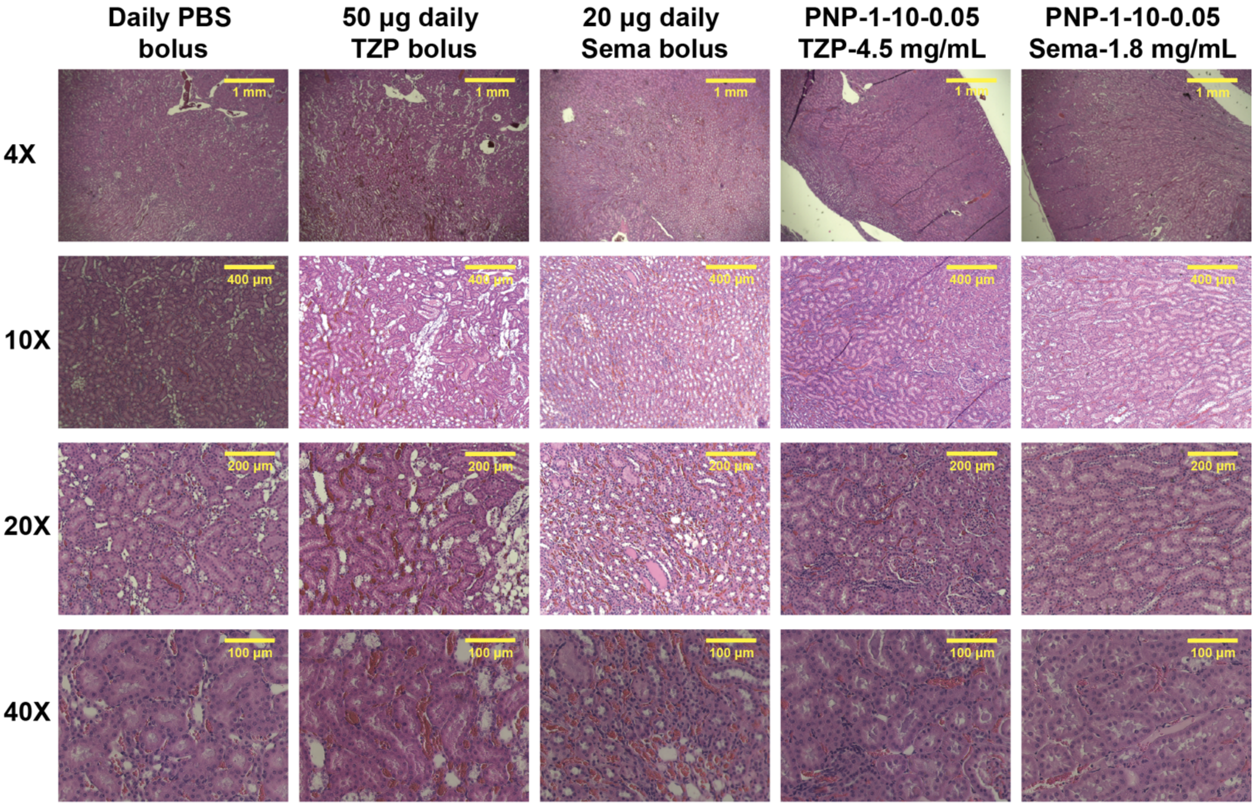

**Figure S4. Liver histology on Day 43.** Representative hematoxylin and eosin (H&E)-stained cross-sections of liver tissue collected 43 days post-treatment. Images illustrate the extent of hepatic cellular integrity, inflammatory infiltration, and tissue architecture across treatment groups.

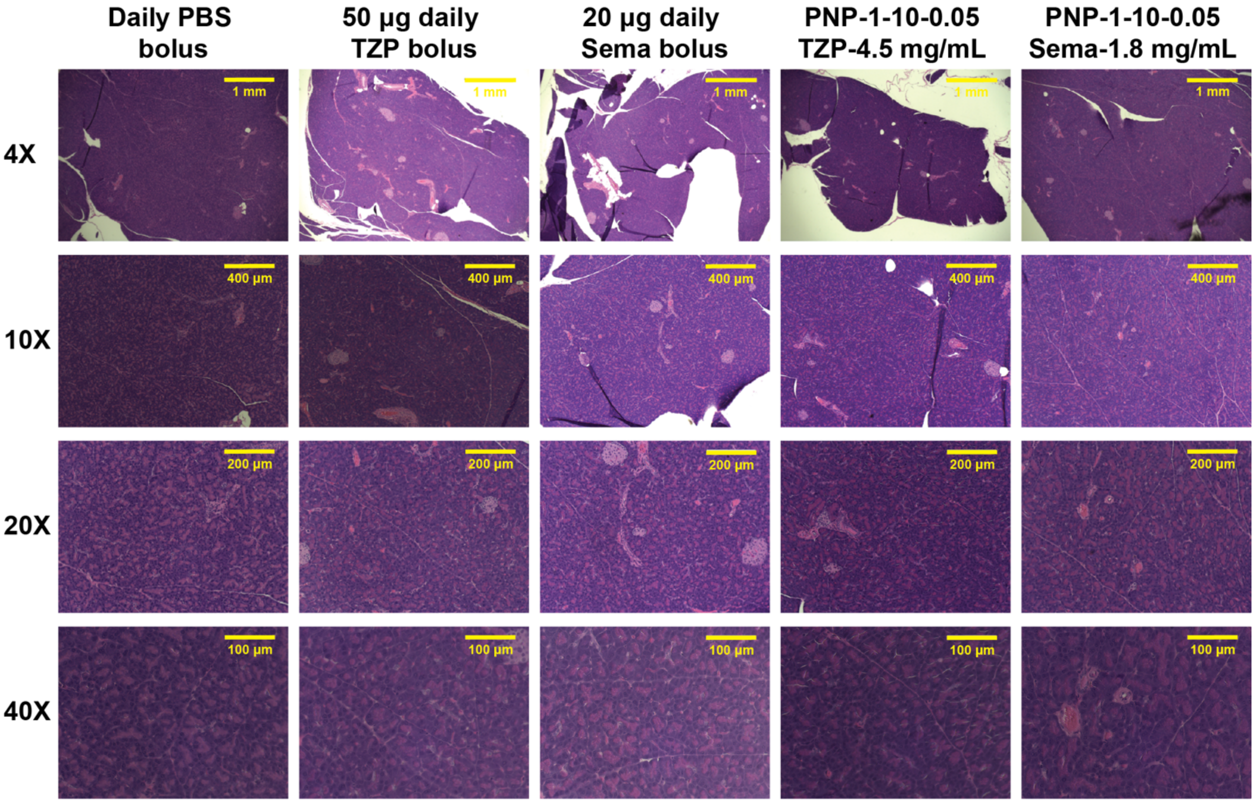

**Figure S5. Pancreas Histology – Day 43.** Representative hematoxylin and eosin (H&E) stained cross-sections of pancreas tissue collected on Day 43 post-treatment. Images depict pancreatic morphology across treatment groups.

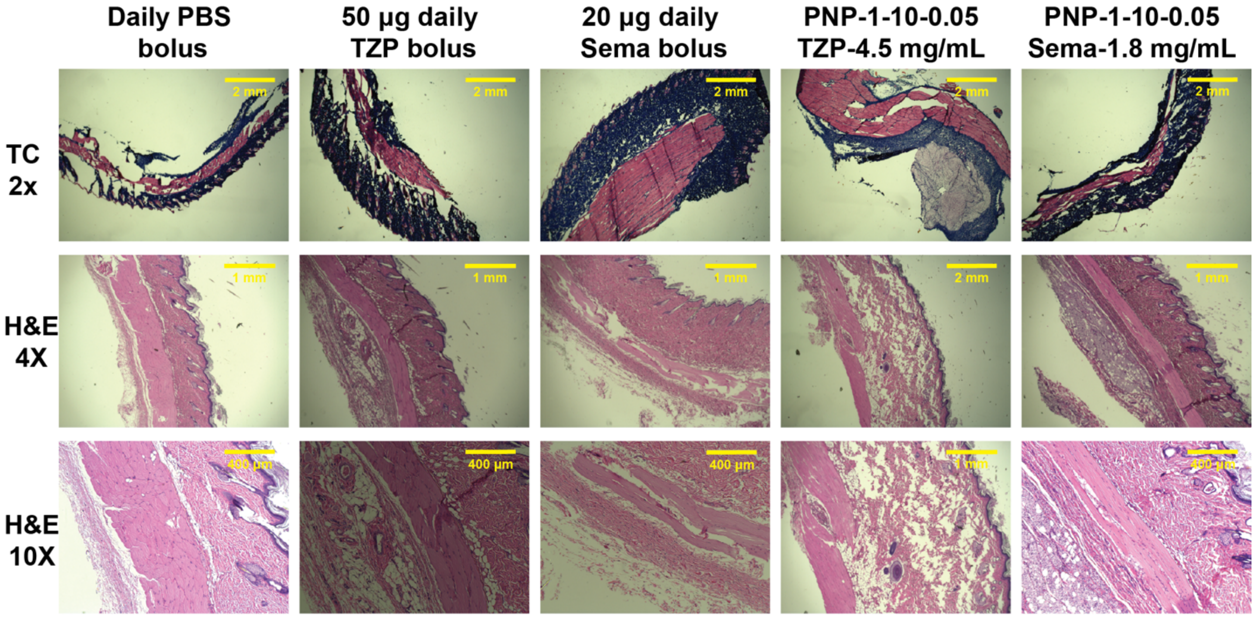

**Figure S6. Skin Histology at Day 43.** Representative cross-sections of skin collected 43 days post-treatment. The top row shows Masson's trichrome-stained samples, highlighting collagen structure and potential fibrosis. The bottom three rows display hematoxylin and eosin (H&E)-stained sections used to assess overall tissue morphology, including epidermal integrity, dermal architecture, and inflammatory cell infiltration.

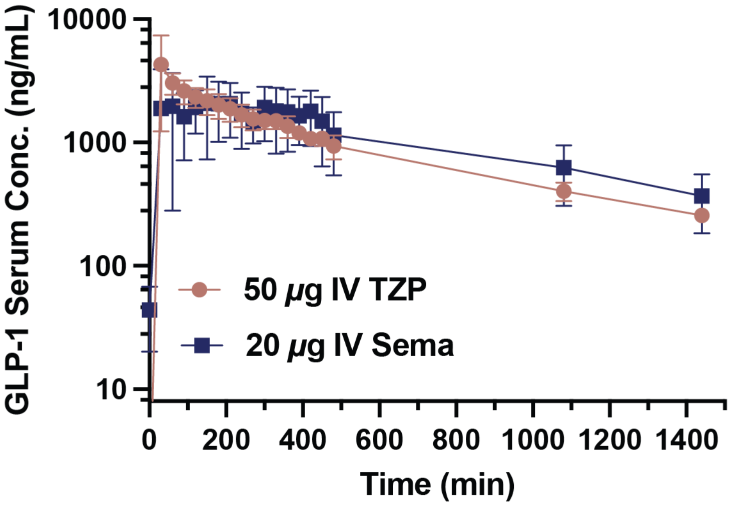

**Figure S7. 24-hour pharmacokinetics in a T2D rat model.** PK profile for Sema (50 μg and 20 μg) TZP administered intravenously as a soluble solution in PBS buffer.

**Table S1. Pharmacokinetic parameters of GLP-1 RAs in rats, IV PK data.**

| GLP-1 RA | Elimination half-life (hours) |
|---|---|
| Semaglutide | 4.59 ± 1.02 |
| Tirzapetide | 3.87 ± 0.63 |

**Table S2. Pharmacokinetic parameters of GLP-1 RAs in rats, SC gel data.**

| GLP-1 RA | Elimination half-life (days) | $C_{max}$ (ng/mL) | AUC (ng/mL) |
|---|---|---|---|
| Semaglutide | 0.71 ± 0.001 | 2213.5 ± 286.2 | 3708.6 ± 1096.6 |
| Tirzapetide | 0.71 ± 0.003 | 3518.3 ± 1175.7 | 3766.6 ± 1426.4 |

**References**

1. Chen, T., L. Kagan, and D.E. Mager, *Population Pharmacodynamic Modeling of Exenatide After 2-Week Treatment in STZ/NA Diabetic Rats.* Journal of Pharmaceutical Sciences, 2013. **102**(10): p. 3844-3851.
